# Bayesian Optimization of Neurostimulation (BOONStim)

**DOI:** 10.1101/2024.03.08.584169

**Authors:** Lindsay D. Oliver, Jerrold Jeyachandra, Erin W. Dickie, Colin Hawco, Salim Mansour, Stephanie M. Hare, Robert W. Buchanan, Anil K. Malhotra, Daniel M. Blumberger, Zhi-De Deng, Aristotle N. Voineskos

## Abstract

**Background:** Transcranial magnetic stimulation (TMS) treatment response is influenced by individual variability in brain structure and function. Sophisticated, user-friendly approaches, incorporating both established functional magnetic resonance imaging (fMRI) and TMS simulation tools, to identify TMS targets are needed.

**Objective:** The current study presents the development and validation of the Bayesian Optimization of Neuro-Stimulation (BOONStim) pipeline.

**Methods:** BOONStim uses Bayesian optimization for individualized TMS targeting, automating interoperability between surface-based fMRI analytic tools and TMS electric field modeling. BOONStim’s Bayesian optimization performance was evaluated in a sample dataset (N=10) using standard circular and functional connectivity-defined targets, and compared to densely sampled grid optimization.

**Results:** Bayesian optimization converged to similar levels of total electric field stimulation across targets in under 30 iterations, converging within 5% error of the maxima detected by grid optimization, and requiring less time.

**Conclusions:** BOONStim is a scalable and configurable user-friendly pipeline for individualized TMS targeting with quick turnaround.

## Introduction

Clinical and experimental transcranial magnetic stimulation (TMS) to brain regions exhibits large inter-individual variation in efficacy [1]. Individuals have unique functional connectivity and brain topography patterns [2–4] and individualized functional magnetic resonance imaging (fMRI) approaches and surface-based brain parcellations offer stronger brain-behavior links as compared to standard group-based parcellations [2,5,6]. Further, treatment response to TMS may be influenced by individual differences in functional brain organization and personalized functional targeting may improve outcomes [7–9].

TMS localization and intensity are also influenced by individual variability in cortical geometry [10], which can be accounted for via electric field simulations [11]. Several optimization approaches have been proposed for maximizing the electric field over cortical regions of interest (ROIs) on an individual level [12–14]. However, the current fast optimization tools only allow for binarized, circular ROIs, despite investigators increasingly using fMRI to define anatomically meaningful non-circular parcels or weighted maps derived from functional connectivity as potential TMS targets. Currently available tools also require in-house software for automation and/or data preprocessing, which may lead to reproducibility and scalability issues [15]. There is a need for more sophisticated, user-friendly approaches incorporating fMRI to identify personalized TMS targets with electric field modeling to account for structural variability and optimize dose delivery [16,17].

Accordingly, we have developed the **B**ayesian **O**ptimization **o**f **N**euro-**Stim**ulation (BOONStim) pipeline. BOONStim is an automated end-to-end pipeline for generating TMS coil positions. Personalized cortical targets are identified via surface-based fMRI analytic tools and TMS electric field modeling is used to optimize coil position for target engagement. BOONStim introduces a novel TMS Bayesian optimization algorithm that allows for more general ROI target definitions and quick turnaround. Here, we present this pipeline and validation analyses.

## Materials and Methods

### Pipeline Overview

BOONStim is available as open-source code (https://github.com/TIGRLab/BOONStim) and an overview is shown in Figure 1. The pipeline supports a modular, parallelizable, and customizable workflow scripted using Nextflow [18]. BOONStim leverages portable, isolated Singularity containers to encapsulate its environments, including widely established fMRI software packages, reducing the burden of computational set-up for users. This also allows users to easily swap out container versions and reduce dependency conflicts between software. A small utility python library (Fieldopt) is packaged with BOONStim that contains the optimization logic (https://github.com/TIGRLab/fieldopt).

**Figure 1.**
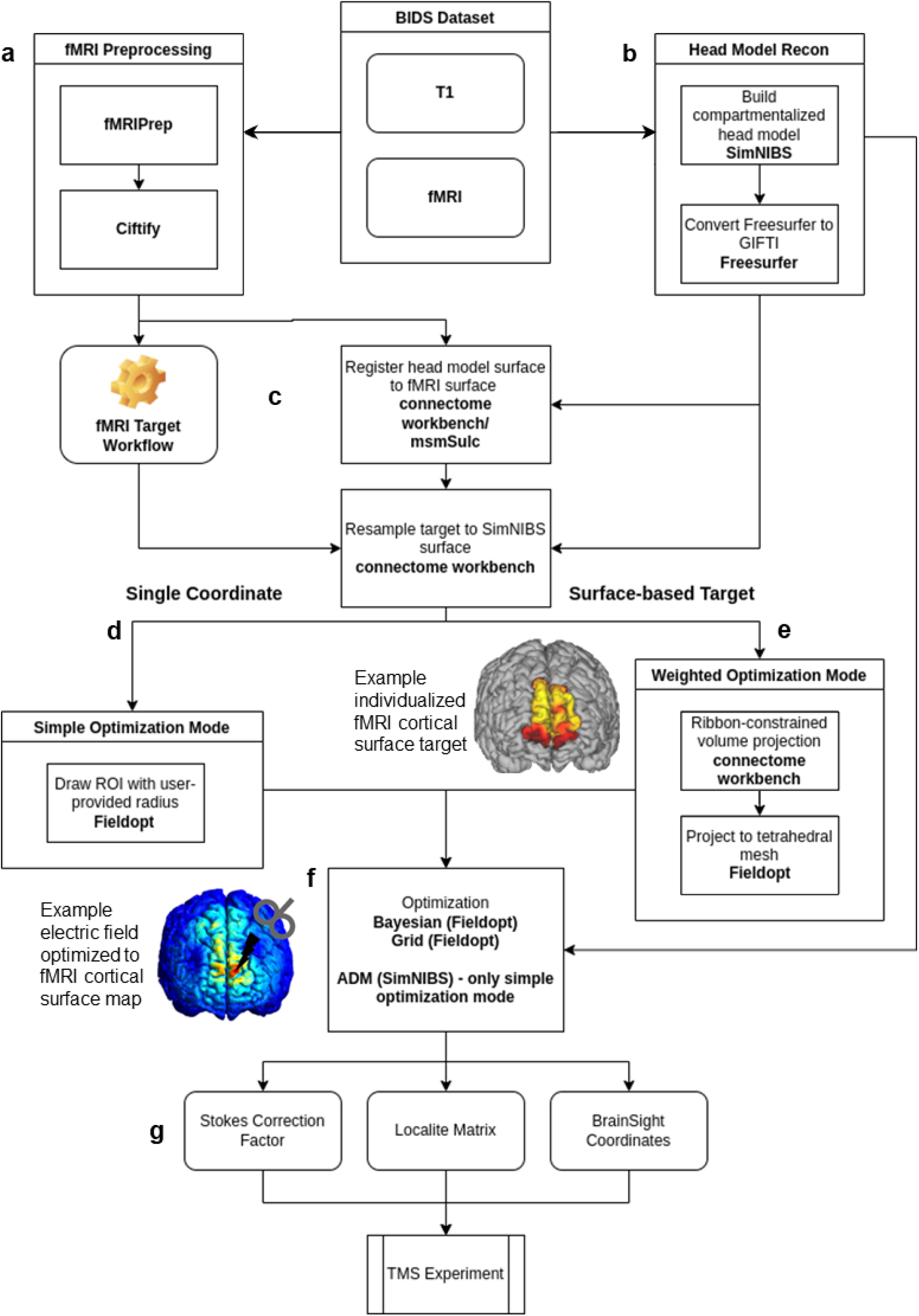
BOONStim pipeline overview BOONStim utilizes Singularity container image files for widely established functional magnetic resonance imaging (fMRI) software packages including fMRIPrep [20], Ciftify [21], HCP (Human Connectome Project) Pipelines [26], SimNIBS (Simulation of Non-invasive Brain Stimulation) [13,14], Connectome Workbench [27], and Freesurfer [22], as well as Fieldopt. Input T1-weighted and fMRI data are required to be organized according to the Brain Imaging Data Structure (BIDS) standard [19], enabling flexible use of established preprocessing tools. **a)** MRI preprocessing is done using fMRIPrep [20], followed by Ciftify [21] to project the data to the cortical surface. **b)** T1-weighted head model reconstruction (recon) is achieved using SimNIBS [13,14] and Freesurfer [22] to generate a finite element method (FEM) model, which is also transformed to the surface (GIFTI format). **c)** Connectome Workbench/MSMSulc [28] is then used to register the head model to the fMRI surface. Users may define their own fMRI target workflows to derive ROIs using available parcellations or outputs from fMRIPrep and Ciftify, or a standard TMS target (i.e., MNI coordinate), after which the target ROI is resampled to the SimNIBS surface using Connectome Workbench [27]. **d)** If the user is targeting a single coordinate, simple optimization mode is used to draw an ROI with the provided parameters using Fieldopt. **e)** If a surface-based target is provided (e.g., functional connectivity maps), weighted optimization mode is used. BOONStim automates interoperation of the user-defined ROI with SimNIBS simulation software by mapping the user ROI directly into the FEM in which simulations are performed. This is achieved via ribbon-constrained volume projection using Connectome Workbench and a tetrahedral projection algorithm using monte carlo sampling to determine contributions of voxels to specific tetrahedra in Fieldopt. **f)** Once the target is represented on the FEM, the choice of optimization is flexible via a single configuration option, including Bayesian optimization or grid optimization. For a single MNI coordinate, there is also the option of using auxiliary dipole method (ADM) optimization [12]. **g)** BOONStim output files include coordinates calculated during optimization and the stokes correction factor, which are ready to use with TMS neuronavigation software and hardware.

BOONStim requires input T1-weighted and fMRI data to be organized according to the BIDS standard [19]. MRI preprocessing is done using fMRIPrep [20] and Ciftify [21], and SimNIBS [13,14] and Freesurfer [22] are used for head model reconstruction, which is registered to the cortical surface. These preprocessing pipelines can be configured to suit various MRI acquisition parameters using boutiques JSONs [23], enabling versioning and reproducibility.

Users may define their own fMRI target workflows to derive ROIs or use a MNI coordinate, which is resampled to the SimNIBS surface. If the user is targeting a single coordinate, an ROI is drawn, whereas a surface-based target requires weighted optimization. Once the target is represented on the head model, our Bayesian optimization approach can be applied, or the current gold standard, grid optimization. Auxiliary dipole method (ADM) optimization can also be used for a MNI coordinate [12].

During optimization, electric field modeling is performed at different locations around the target. Unlike grid optimization, which tests every unique combination of coil angles and positions in a grid around the target, Bayesian optimization learns from tested parameter combinations to inform the next iteration, reducing dependence on the search space size and computational cost. BOONStim’s Bayesian optimization process learns from a history of sampling points with basic assumptions about the smoothness of the underlying objective function, minimizing the number of iterations to find a global maximum. It uses the information criterion called “multipoints expected improvement” or the q-EI, enabling parallel sampling for faster convergence [24]. The optimization objective is defined as follows:

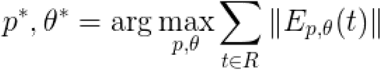

Where

*p* - a position vector specifying the centre of the TMS coil

*θ -* the orientation of the TMS coil handle

*t* - a tetrahedron from the head model

*E*_*P,θ* (*t*)_ *-* the electric field vector at a given tetrahedron *t* from a simulation of a coil at (*p. t*)

*R* - a region of interest

BOONStim produces outputs which are directly usable with established TMS neuronavigation software, including BrainSight and Localite inputs from the coordinates calculated during optimization, and the stokes correction factor based on an automated scalp-to-cortex distance assessment for both hemispheres [25]. It also provides a reference image, so the TMS operator can visually confirm the coil position, and the corrected T1 image to ensure the same coordinate system is used. Quality control pages are generated for each subject with results from sub-workflows (i.e., fMRIPrep, Ciftify, and SimNIBS).

### Dataset

BOONStim was run on 10 subjects from the Dual Mechanisms of Cognitive Control dataset, intended for comparing preprocessing pipelines [29]. Data was acquired using a 3T Siemens Prisma Fit scanner, including an anatomical T1-weighted scan (isotropic resolution=0.8 mm, TR=2400 ms), two resting-state fMRI scans with opposite phase encoding directions (isotropic resolution=2.4 mm, TR=1200 ms, duration=5.2 minutes per scan), and field map scans. See OpenNeuro Dataset: ds003452 for details.

The performance of BOONStim’s Bayesian optimization was evaluated using standard circular ROIs of the dorsolateral prefrontal cortex (DLPFC), the most common TMS target, and functional connectivity maps of the dorsomedial PFC (DMPFC), a core node in social cognitive processing [30], because Boonstim was developed as part of a clinical trial targeting this region. Bayesian optimization was compared to densely sampled grid optimization.

### Target Definitions

Standard circular ROI targets were produced by drawing a circle (5, 10, and 20 mm radius) around the selected vertex (MNI coordinates:-46,45,38 [31]). Masked functional connectivity maps were used to generate functional connectivity-defined DMPFC targets. The DMPFC region was removed from a “mentalizing” map from Neurosynth [32], which was projected onto the cortical surface and used to compute mentalizing seed-based functional connectivity for each participant. A binarized DMPFC mask was applied to the seed-based functional connectivity map to produce personalized DMPFC-mentalizing connectivity maps to optimize over.

### Bayesian Optimization Validation

To verify the results of Bayesian optimization, we compared it to densely sampled grid optimization. For Bayesian optimization, we performed 30 optimizations for each participant given that it is stochastic. Grid optimization was performed for each participant using a 30x30 grid of locations with 36 orientations (5 degree increments), resulting in 32400 unique coil positions being evaluated.^1^ The coil position with a maximal dosage (i.e., simulated electric field stimulation) over the target was selected as the optimal position for a given participant and ROI.

For both targets, we computed the number of iterations required (up to a maximum of 30) for Bayesian optimization to converge onto the ‘gold standard’ maxima detected by a densely sampled grid optimization, the percent difference between the simulated electric field stimulation over the target region using grid optimization and Bayesian optimization for each participant, and the number of iterations required to reach convergence.

## Results

For both the standard circular ROI targets (Figure 2a) and the functional connectivity-based targets (Figure 2b), Bayesian optimization converged to similar levels of total electric field stimulation compared to grid optimization across participants in under 30 iterations.

**Figure 2.**
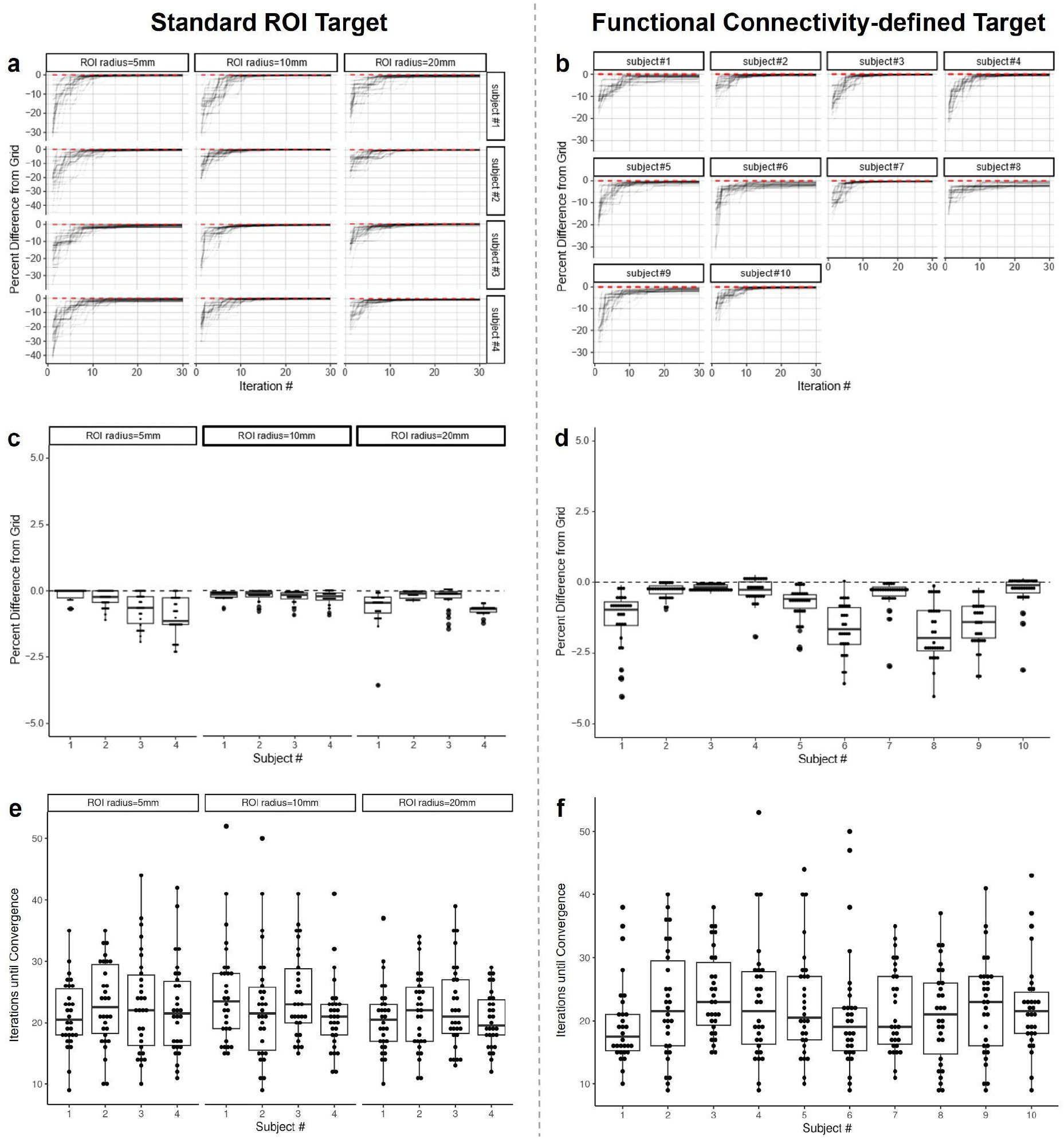
Bayesian Optimization Validation **Left** panels display results for optimizing over the standard ROI target (DLPFC) with a radius of 5 mm, 10 mm, and 20 mm for four representative subjects, and **right** panels for optimizing over the functional connectivity-defined target (DMPFC) for all 10 subjects. **a-b** display the percent difference of Bayesian optimization from grid-based optimization maxima as the number of iterations increases. Percent differences are plotted because the objective value achieved by the optimization varies across subjects based on the size of the ROI, scalp-to-cortex distance, and functional connectivity. Each line represents one of 30 optimizations performed for each participant, and the red dotted line at 0 represents the maxima detected by densely sampled grid optimization. **c-d** show the percent difference of Bayesian optimization from grid-based optimization maxima, where each data point represents one of 30 optimizations performed for each participant. **e-f** display the number of iterations required for Bayesian optimization to reach convergence, where each data point represents one of 30 optimizations performed for each participant.

Bayesian optimization converged to <5% error of the maxima detected by grid optimization (Figure 2c and 2d), performing similarly across standard ROI targets of different sizes and functional connectivity-defined targets. The total number of iterations required to achieve convergence ranged from 9 to 52 across participants for standard ROI targets (mean(SD)=22(6.9); Figure 2e) and from 9 to 53 for functional connectivity-defined targets (mean(SD)=22(8.0); Figure 2f), compared to 32400 iterations for densely sampled grid optimization.

## Discussion

BOONStim represents a scalable end-to-end pipeline that facilitates individualized fMRI-guided TMS targeting. BOONStim incorporates a novel Bayesian optimization algorithm that solves TMS optimization over generalized ROIs, performing similarly to densely sampled grid-based optimization with low percentage error and a substantially reduced run-time. Three key advantages that BOONStim has over other methods include:

### 1. Efficiency

Bayesian optimization reached convergence in far fewer iterations than grid-based optimization for both standard and functional connectivity-defined ROIs, requiring much less time than dense sampling. Its relative invariance to search space size without sacrificing performance is significant in the context of clinical trials where turnaround times between acquiring fMRI data and delivering TMS treatment can be critical.

### 2. Generality across target ROIs

BOONStim can use weighted scalar-valued functional connectivity maps as inputs, and its Bayesian optimization was not significantly impacted in its ability to converge to a maxima across various sizes, shapes, and types of ROIs across participants. This approach is successfully being used in ongoing clinical trials targeting both the DMPFC (NCT06118268) and the DLPFC (NCT05583747).

### 3. User-friendly interoperability of established fMRI and TMS tools

BOONstim integrates the current best fMRI data processing practices and automates operability with TMS simulation software. Its scalability allows researchers to run high-throughput TMS experiments without worrying about many pipeline implementation details.

## Conclusions

BOONStim enables researchers to perform automated individualized coil optimization in TMS trials with minimal setup and define rich targets with variable shapes and weights, such as functional connectivity maps. BOONStim is a tool built on reproducible fMRI analytic approaches that serves as a scalable and configurable end-to-end solution for TMS practitioners to reduce the burden of performing sophisticated fMRI-based targeting with electric field modeling. This pipeline is also extensible to newer optimization techniques and fMRI methodology.

## CRediT authorship contribution statement

**Lindsay D. Oliver**: Writing - Original Draft, Review & Editing, Formal Analysis, Visualization. **Jerrold Jeyachandra**: Conceptualization, Methodology, Software, Formal Analysis, Visualization, Writing - Original Draft Preparation, Review & Editing. **Erin W. Dickie**: Conceptualization, Methodology, Writing - Review & Editing. **Colin Hawco**: Conceptualization, Methodology, Writing - Review & Editing. **Salim Mansour**: Software, Formal Analysis, Writing - Review & Editing. **Stephanie M. Hare**: Writing - Review & Editing. **Robert W. Buchanan**: Funding Acquisition, Writing - Review & Editing. **Anil K. Malhotra**: Funding Acquisition, Writing - Review & Editing. **Daniel M. Blumberger**: Funding Acquisition, Methodology, Writing - Review & Editing. **Zhi-De Deng**: Methodology, Software, Supervision, Writing - Review & Editing. **Aristotle N. Voineskos**: Conceptualization, Supervision, Funding Acquisition, Writing - Original Draft, Review & Editing.

## Declarations of interest

The authors report no financial relationships with competing interests.

## Funding/Support

This work was supported by the National Institute of Mental Health grant R61 MH120188 (to ANV and DMB) and the Brain & Behavior Research Foundation Young Investigator Grant (to LDO).

## Role of the Funder/Sponsor

The funding organizations had no role in the design and conduct of the study; collection, management, analysis, and interpretation of the data; preparation, review, or approval of the manuscript; and decision to submit the manuscript for publication.

## Footnotes

1 It should be noted that this dense sampling space is not the default if grid optimization is selected in BOONStim, but grid density parameters are configurable.

## Notes

### Competing Interest Statement

The authors have declared no competing interest.

### Summary of Updates

We have updated the manuscript and authorship information to reflect additional feedback and revisions.

